# Convergent patterns of sex chromosome dosage compensation between lepidopteran species (WZ/ZZ) and eutherian mammals (XX/XY): insights from a moth neo-Z chromosome

**DOI:** 10.1101/030767

**Authors:** Liuqi Gu, James R. Walters, Douglas C. Knipple

## Abstract

In contrast to XX/XY species, Z-linked expression is overall reduced in female WZ/ZZ species compared to males or the autosomal expression. This pattern (Z<ZZ≈AA) has been consistently reported in all the WZ/ZZ taxa examined so far, with the singular exception of the insect order of Lepidoptera (moths and butterflies). However, conflicting results linger in this taxon due to discrepancies in data analyses and tissues sampled. To address this issue, we analyzed dosage compensation in the codling moth *Cydia pomonella* (Tortricidae) using tissues that represent different levels of sexual divergence. *C. pomonella* is the most basal lepidopteran species yet examined for dosage compensation and has a neo-Z chromosome resulting from an ancient Z:autosome translocation. We based our analyses on RNAseq and *de novo* transcriptome data from *C. pomonella,* as well as scrutiny into investigations of other lepidopteran species. Our evidence supports that the lepidopterans share a pattern (Z ≈ ZZ < AA) of dosage compensation that mirrors the eutherian mammals (X ≈ XX < AA). In particular, reproductive tissues appear to be exempt from dosage compensation, which helps explain the incongruence in prior reports. Furthermore, *C. pomonella* ancestral-Z segment exhibited a greater expression reduction than genes on the neo-Z segment, which intriguingly also reflects the differential up-regulation between the ancestral and newly-acquired X-linked genes in mammals. The insect order of Lepidoptera challenges both the classic theories regarding evolution of sex chromosome dosage compensation and the emerging view on dosage compensation’s association with sexual heterogamety.

## Introduction

In the animal kingdom, there are two predominant sex chromosome constitutions with opposing patterns. In XX/XY species, females are the homogametic sex (XX) and males are the heterogametic sex (XY). This pattern is reversed in WZ/ZZ species, where females are heterogametic (WZ) and males are homogametic (ZZ). In both cases, however, sex chromosome dosage compensation is expected to evolve concomitantly with the sex chromosomes to 1) equalize sex-linked gene expression between the male and female (X~XX | Z~ZZ compensation) and 2) balance the gene expression from the sex chromosome with that from the autosomes in the heterogametic sex (X~AA | Z~AA compensation). Different organisms deploy distinct strategies to achieve the two aspects of dosage compensation. In XX/XY systems, the X~XX compensation involves orchestrated chromosome-wide regulation (reviewed in (Ferrari, et al. 2014)). In *Drosophila,* transcriptional doubling of the single X chromosome of XY males equalizes its output both to that from two X copies of XX females. This male X hyper-expression also balances expression relative to the autosomes (AA) (i.e., X≈XX≈AA, reviewed in (Conrad and Akhtar 2012; Gelbart and Kuroda 2009; Laverty, et al. 2010)). In contrast, in both placental mammals and *Caenorhabditis* worms, X~XX compensation is achieved by reducing X expression in the homogametic sex. Mammals inactivate one X chromosome (reviewed in (Schulz and Heard 2013)) while worms repress both X chromosomes by half (reviewed in (Meyer 2000, 2010)), respectively. Additionally, general two-fold up-regulation (global X~AA compensation), as first proposed by Ohno (Ohno 1967), is expected to match the virtually monoallelic X expression in both sexes to the biallelic autosomal expression (i.e., X≈XX≈AA). However, this theory is widely challenged and growing evidence favors the reduction in X expression in both mammals (reviewed in (Pessia, et al. 2014)) and *C. elegans* (Albritton, et al. 2014) (i.e., X≈XX<AA).

Patterns of dosage compensation in WZ/ZZ species generally have shown a pattern distinct from XX/XY systems, particularly among vertebrates. Female heterogametic species typically lack global Z~ZZ compensation and therefore exhibit Female:Male (F:M) expression disparity on the Z chromosome. This Z<ZZ≈AA pattern has been reported in a wide range of taxa with independently-evolved female heterogamety, including birds (Adolfsson and Ellegren 2013; Ellegren, et al. 2007; Itoh, et al. 2007; Itoh, et al. 2010; Uebbing, et al. 2013; Wolf and Bryk 2011), *Schistosoma mansoni* (a trematode parasite) (Vicoso and Bachtrog 2011), snakes (Vicoso, et al. 2013) and *Cynoglossus semilaevis* (a flatfish) (Chen, et al. 2014). To date, the only exceptions to this pattern come from the insect order of Lepidoptera (moths and butterflies), but even among Lepidoptera the results are varied. Although the same pattern of Z<ZZ≈AA was first reported in the silk moth *Bombyx mori* (Zha, et al. 2009), this result was later deemed problematic due to artifacts of microarray normalization and a failure to examine F:M expression ratio on autosomes along with the Z chromosome (Walters and Hardcastle 2011). Re-analyses of the same data indicated a Z≈ZZ<AA pattern, which presents a surprising analogue to that found in mammals. However, doubts about this counterintuitive result lingered due to the robustness in the microarray data itself (Harrison, et al. 2012). Using RNAseq data, Harrison et al. (2012) reported a Z<ZZ≈AA pattern in the Indian meal moth *Plodia interpunctella,* which further strengthened the popular speculation that F:M disparity in sex-linked expression is universally associated with female heterogamety (Graves and Disteche 2007; Harrison, et al. 2012; Mank 2013, 2009; Naurin, et al. 2010; Vicoso and Bachtrog 2009; Wolf and Bryk 2011). In contrast, however, two more recent RNA-seq studies of *Manduca sexta* (Smith, et al. 2014) and *Heliconius spp.* (Walters, et al. 2015) also revealed the Z≈ZZ<AA pattern, in good agreement with the re-analyses of the *B. mori* microarray data by Walters et al (2011).

Unfortunately, these studies differed in the nature of transcriptome data (microarray versus RNAseq) and tissues sampled (Fig 1). In particular, there has been no RNAseq data specifically for reproductive tissues, while the *P. interpunctella* study used whole-body adult insects, which contains a substantial fraction of reproductive tissues. Ideally, gonads are isolated in dosage compensation studies because of the widely-observed germline-specific regulation of the sex chromosomes, such as absence of dosage compensation (Kelly, et al. 2002; Meiklejohn, et al. 2011; Sugimoto and Abe 2007) or meiotic sex chromosome inactivation (Bean, et al. 2004; Guioli, et al. 2012; Monesi 1965; Schoenmakers, et al. 2009; Turner 2007; Vibranovski 2014). In addition, genes with predominant expression in gonads are often disproportionally represented on the sex chromosomes compared to the autosomes (Arunkumar, et al. 2009; Khil, et al. 2004; Lercher, et al. 2003; Parisi, et al. 2003; Reinke, et al. 2004; Saifi and Chandra 1999). All these factors add layers of complexity to the assessment of dosage compensation and potentially bias the results when gonads and soma are analyzed together.

**Fig 1.**
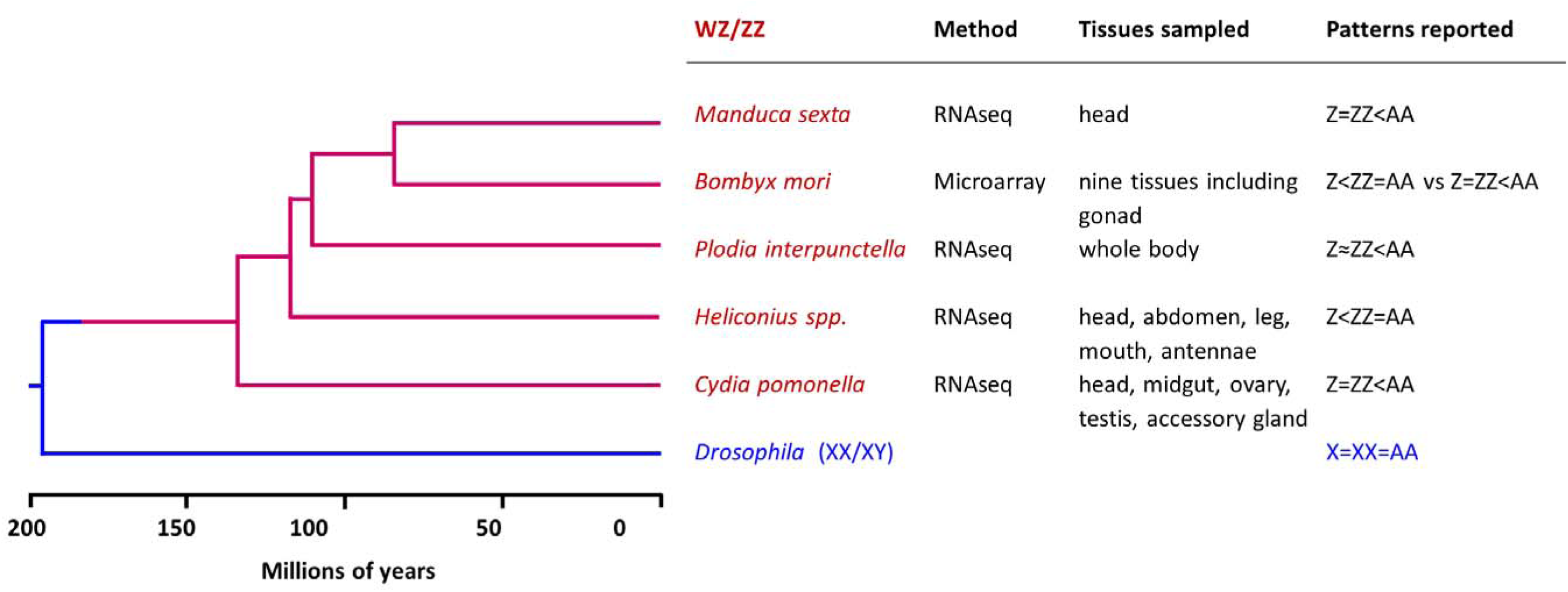
Summary of dosage compensation studies in Lepidoptera. Phylogenetic relationship among insect species mentioned in this study is based on (Marec, et al. 2010; Wahlberg, et al. 2013).

In order to gain further insights on dosage compensation in the Lepidoptera, we generated RNA-seq data from the codling moth *Cydia pomonella* and performed comprehensive analyses that permit comparisons with previous studies. *C. pomonella* represents a more basal phylogenetic group (Tortricidae) compared to the other lepidopterans previously examined for dosage compensation (Fig 1). More importantly, *C. pomonella* has a neo-Z chromosomal segment that reflects an autosomal fusion with the ancestral Z chromosome (Fig 2) (Nguyen, et al. 2013). Both molecular and cytogenetic evidence showed that *C. pomonella* W chromosome is extensively degenerated as in other Lepidopteran species (Fukova, et al. 2007; Nguyen, et al. 2013; Sichova, et al. 2013). Therefore, the autosomal progenitors of *C. pomonella* neo-Z genes were brought under the same sex-linked inheritance and accompanying sex-specific gene dosage as the ancestral-Z (ancl-Z) genes. This evolutionary partitioning of the *C. pomonella* Z chromosome provides a novel opportunity to examine the effect of sex-linkage on dosage compensation.

**Fig 2.**
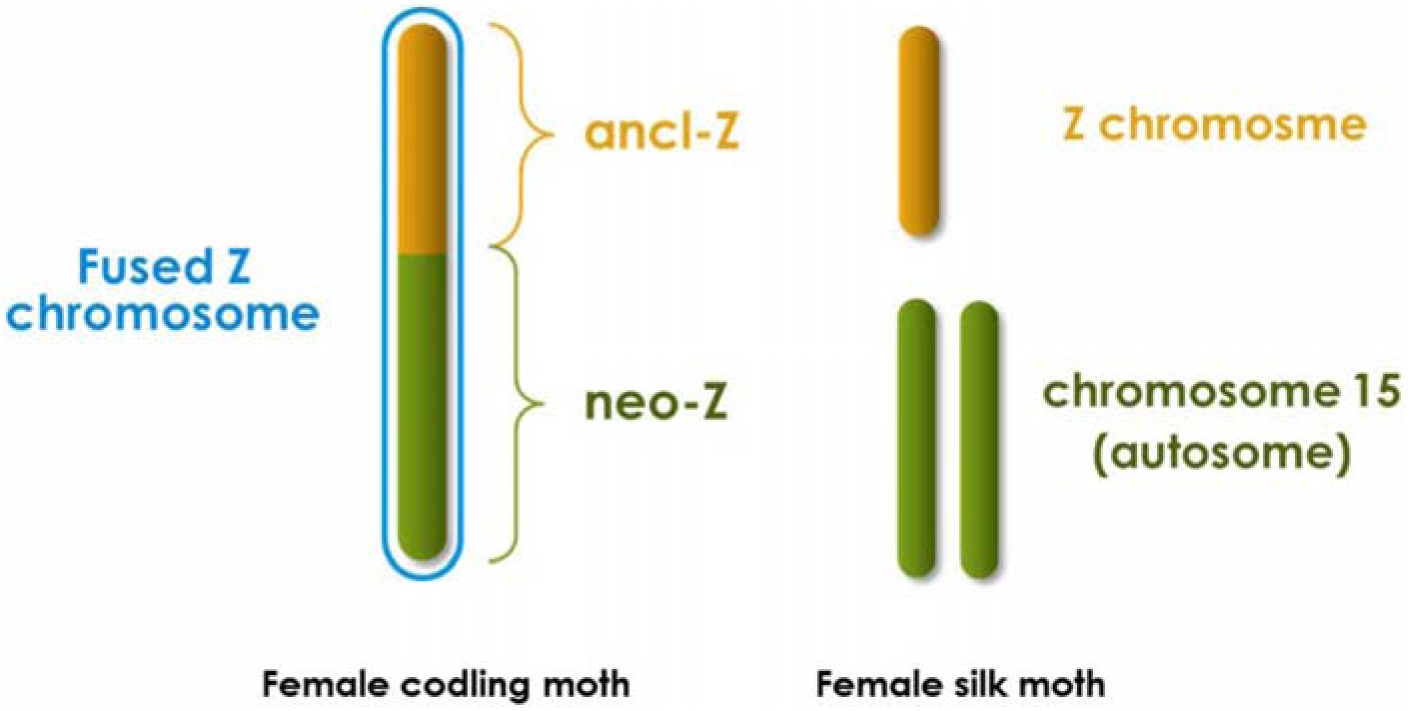
Sex chromosomes comparison between *C. pomonella* and *B. mori.* The ‘ancl-Z’ and ‘neo-Z’ chromosomal segments of *C. pomonella* are homologous to the Z chromosome and autosome 15 of *B. mori,* respectively. Z chromosomes are effectively hemizygotic in lepidopteran females; W chromosomes contain no functional genes and are thus not drawn.

## Results and Discussion

### Samples, RNAseq, *de novo* assembly and chromosomal assignments

We sampled seven *C. pomonella* adult female (F) and male (M) tissues that represent different levels of sexual divergence: midgut (F|M), head (F|M), ovary (F), testis (M) and accessory gland (M). The digestive tissue of the adult midgut in the abdomen should have minimal sexual dimorphism since feeding activity is limited during lepidopteran adulthood. Insect adult head is the olfactory center for courtship and mating and is thus presumed to be one of the most sexually dimorphic tissues in the insect soma. Reproductive tissues are sex-specific, representing a maximum of sexual dimorphism. Among the male reproductive tissues (Fig 3), testis is the gonadal counterpart of the ovary in females, whereas the remaining components are of somatic origin (sex-specific somatic tissues). The male reproductive tract was sampled separately from the testis and simply treated as a single tissue, referred to as ‘accessory gland’ in this analysis.

**Fig 3.**
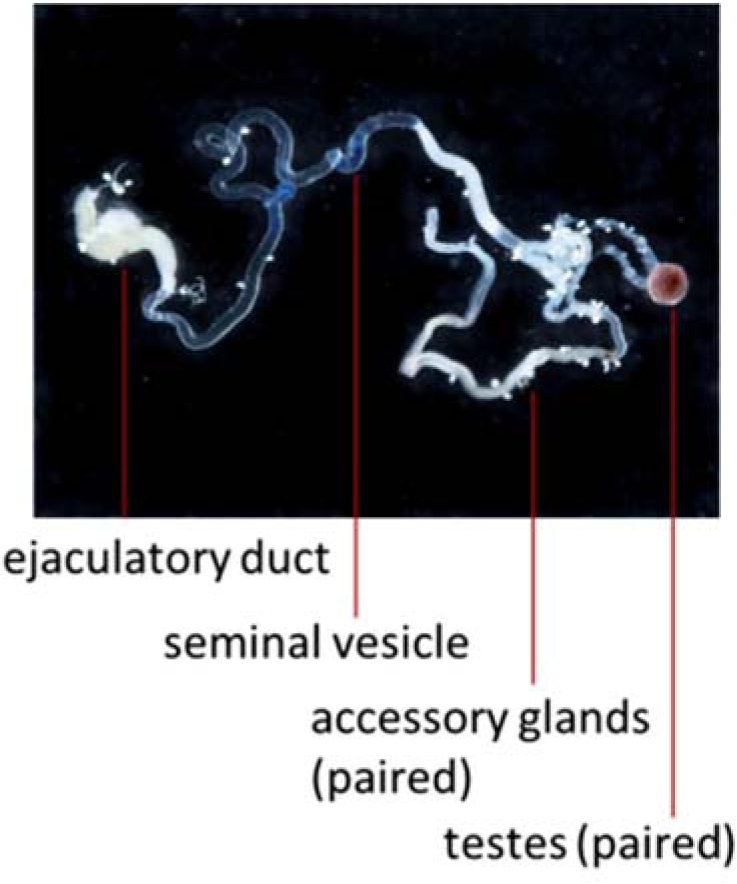
Male reproductive tissues of *C. pomonella.* Accessory gland (the predominant component), along with seminal vesicles, ejaculatory duct and ejaculatory bulb (not shown) comprise the adult male reproductive tract. We collected these male reproductive components, which are of somatic origin, as a single sample and simply call it ‘accessory gland’.

We constructed a *de novo* assembly from the combined transcriptome of these *C. pomonella* tissues, using 238 million pairs of post-filtering reads yielded from RNA sequencing. Conserved synteny in the Lepidoptera allows us to assign putative chromosomal locations of *C. pomonella* genes based on homology to the *B. mori* reference genome. Among assembled contigs, we identified 447 *C. pomenella* genes to be putatively located on the ancestral Z segment (ancl-Z), 535 on the neo-Z segment, and 9,053 on the autosomes. Only these 10,616 contigs were used in all subsequent analyses.

Gene expression intensity is commonly measured by normalizing read counts into fragment per kilobase per million mappable reads (FPKM). In addition, we applied a scaling method, Trimmed Mean of *M*-values (TMM) to normalize FPKM values, which enables comparisons among tissue samples that contain distinctive total RNA populations (Robinson and Oshlack 2010).

### Pattern of dosage compensation (Z≈ZZ<AA) in non-reproductive somatic tissues

Using the *de novo* transcriptome contigs assigned to chromosomal classes, we first assessed dosage compensation in non-reproductive somatic tissues: head and midgut. For Z~ZZ compensation, we evaluated F:M expression using all genes co-expressed (FPKM > 0) in both sexes for a given tissue. In both head and midgut, sex parity of expression was observed regardless of chromosomal location, with median F:M ratios for autosomal, ancl-Z and neo-Z genes all close to 1 in both head and midgut (Fig 4A). The distribution of F:M expression ratios did not differ significantly between autosomes and either of the two Z chromosomal segments (p >0.1, Benjamini-Hochberg-corrected Komolgorov-Smirnov test). This lack of gene dosage effect on the Z chromosome indicates that *C. pomonella* completely compensates Z-linked expression for differences in gene dosage between the sexes (i.e. complete Z~ZZ compensation) in non-reproductive somatic tissues.

**Fig 4.**
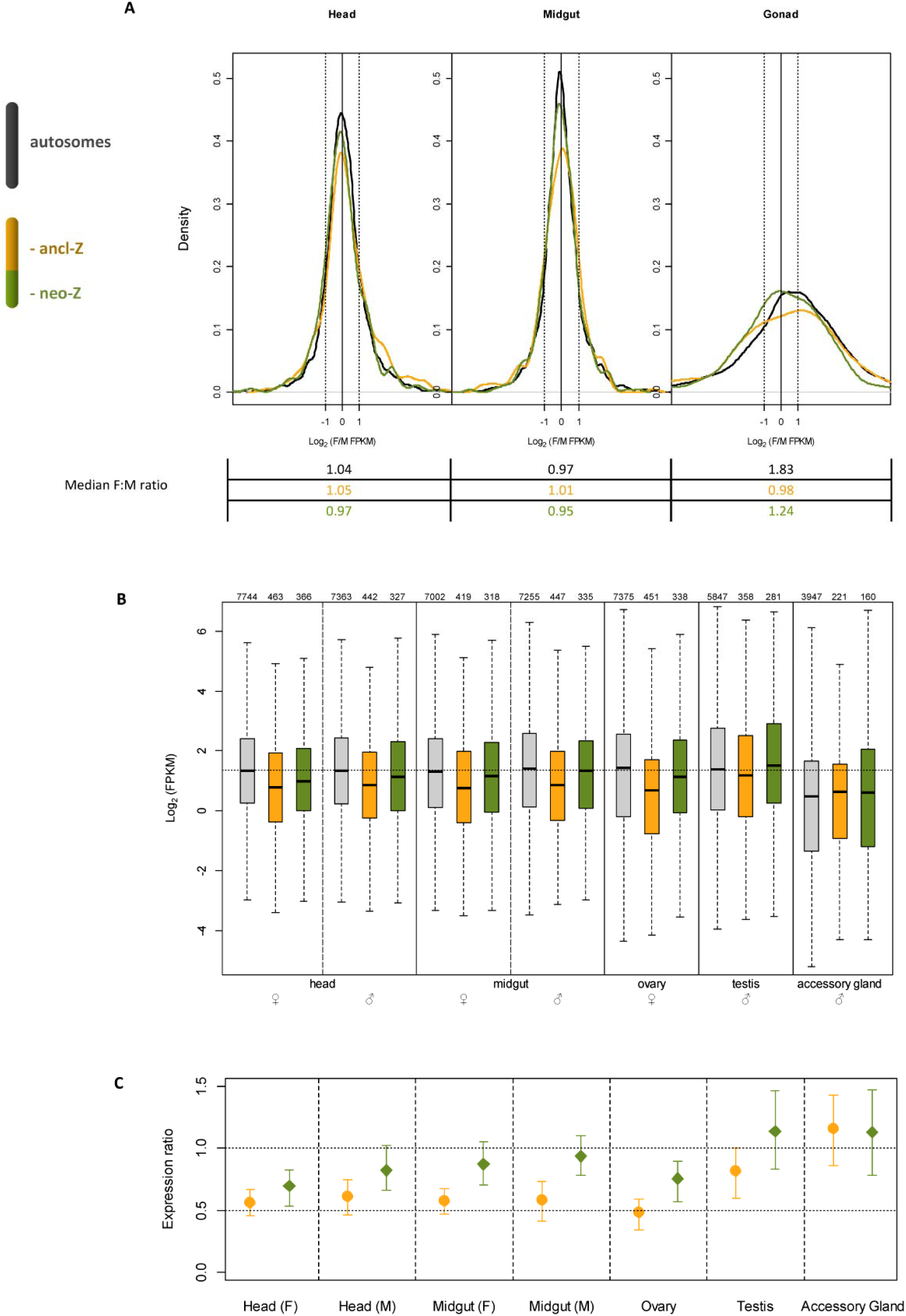
Dosage compensation analysis in *C. pomonella* based on intra-specific data. (A) Z~ZZ compensation in *C. pomonella* tissues assessed by female over male expression (log_2_ transformed FPKM). Key reference log_2_ ratios are marked: -1 (two-fold male over female expression); 0 (equal expression between sexes) and 1 (two fold female over male expression). (B) Expression levels (log_2_ transformed FPKM) of all expressed genes (FPKM>0) in each tissue. The number of total observations is indicated above each boxplot. Plot boxes represent the median and interquartile range of expression levels. The whiskers extend to the most extreme data point that is no more than 1.5 times the interquartile range. The dotted horizontal line across the whole plotting area denotes the mean autosomal expression in the head (F&M) and midgut (F&M). (C) Z(Z):AA ratios of median expression of all expressed genes (FPKM>0) in each tissue. Error bars show 95% confidence intervals estimated by 1000 bootstrap replicates. The horizontal dotted lines denote the reference ratios: 0.5 indicates equal Z(Z) expression to monoallelic autosomal expression (A); 1 indicates equal Z(Z) expression to biallelic autosomal expression (AA). The ancl-ZZ:AA ratios were 0.82 and 1.15 in testis and accessory gland, while the ancl-Z(Z):AA ratios ranged from 0.48–0.61 in the other tissues. The neo-ZZ:AA ratios were 1.14 and 1.13 in testis and accessory gland, while the neo-Z(Z):AA ratios ranged from 0.7–0.93 in the other tissues.

For ‘Z~A compensation’, we contrasted gene expression between the ancl-Z and neo-Z segments with the autosomes in each sex-specific tissue using all genes expressed (FPKM > 0) in that tissue. In all four tissues (head and midgut, both sexes), ancl-Z expression was significantly reduced relative to autosomes (p < 5 × 10^−5^, Bonferroni-corrected Mann-Whitney U test, Fig 4B, S1 Table), with ancl-Z:AA or ancl-ZZ:AA ratios (collectively symbolized as ancl-Z(Z):AA) not significantly different than 0.5 (Fig 4C, bootstrap tests). This ancl-Z(Z):AA ratio of 0.5 indicates functionally monoallelic expression levels for ancl-Z genes relative to diploid autosomal expression levels, which is consistent with incomplete, or absent of, Z~A compensation. In contrast, neo-Z expression was significantly reduced relative to autosomes in head (p < 0.05 in both sexes) but not in midgut (p > 0.3 in both sexes). Except in head (F), the neo-Z(Z):AA ratios were all significantly greater than 0.5 and not significantly different from 1. Taken together, these observations indicate that in *C. pomonella* head and midgut, the extent of Z~A compensation is greater for neo-Z genes than for ancl-Z genes.

Since differences in data structure may influence median-based estimates, we further examined the distribution of expression levels, which has not been previously performed in any lepidopteran studies. In *C. pomonella* head and midgut (both sexes), almost the entire distributions of expression levels of ancl-Z genes not only shift significantly toward lower values compared to those of autosomal genes (p < 0.0005, Benjamini-Hochberg-corrected Komolgorov-Smirnov test), but also did not differ significantly from the distributions of computationally-halved autosomal expression levels (p > 0.2, Benjamini-Hochberg-corrected Komolgorov-Smirnov test, S2 Fig and S2 Table). Similarly for neo-Z genes, comparisons of expression levels with autosomal genes also well corroborate the median-based assessment. When we extended the same analysis to the *M. sexta* and *H. spp.* (head) data sets, we revealed similar shifts toward lower values in both cases, with tests of significant differences for expression level distributions in good agreement with incomplete Z~A compensation as reported in the two original studies (p<0.01, Komolgorov-Smirnov test, S3 Fig).

In addition to supporting median-based assessment of Z~A compensation, our analyses also yield new insights against arbitrarily setting FPKM filtering thresholds to exclude genes with weak expression (also see (Jue, et al. 2013)). For two data groups with different distributions, removing lower values under a same threshold would naturally compress the median values of both groups and eventually make them statistically indistinguishable. Such a ‘compression effect’ becomes more substantial when the group with the lower median also has a larger proportion of low values. This happens to be the case when comparing Z-linked and autosomal expressions in the lepidopteran data sets, as shown by the distribution of expression levels. As shown in the *M. sexta* study (Smith, et al. 2014), increasing FPKM filtering threshold would diminish the Z(Z):AA disparity observed in the unfiltered data (all FPKM>0 values). Under the filtering threshold of 4, the difference begins to appear insignificant. However, a minimum threshold of 4 is not trivially low compared to the median values (~10 and ~12 for Z and autosomes, respectively) and using that threshold, a substantially larger proportion of Z genes (21.6%) are removed compared to autosomal genes (13.4%) (S3 Fig).

### Comparative transcriptome analyses confirm the equal reduction of Z-linked expression in both sexes in the head of *Cydia pomonella.*

Compared to assessing X~XX or Z~ZZ compensation, which involves the same genes under different ‘conditions’ (in male and female), evaluating X|Z~A compensation is somewhat problematic because it typically involves contrasting the expression between sets of non-homologous genes (i.e., autosomal versus sex-linked). Nevertheless, the essence of X|Z~A compensation is to restore the expression of sex-linked genes to their ancestral levels on the proto-sex chromosomes (Ohno, et al. 1959), which cannot be measured directly but may be inferred by the expression of their autosomal orthologs in related species (Albritton, et al. 2014; He, et al. 2011; Julien, et al. 2012; Lin, et al. 2012; Nozawa, et al. 2014). For *C. pomonella,* orthologs of neo-Z genes are autosomal in most other lepidopterans. We can thus not only confirm but also quantify the expression reduction of *C. pomonella* neo-Z genes in the head using the existing datasets from *M. sexta* (Smith, et al. 2014) and *Heliconius melpomene* (Walters, et al. 2015). Robustness of this comparative transcriptome method was supported by the substantial correlation in expression levels between interspecific orthologous pairs (Pearson’s ρ = 0.55 for the *C. pomonella~M. sexta* pair and ρ = 0.4 for the *C. pomonella~H. melpomene* pair). Orthologs of *C. pomonella* ancl-Z, neo-Z and autosomal genes in the reference species are denoted as 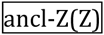, 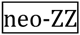 and 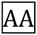, respectively.

Using either *M. sexta* or *H. melpomene* as the reference, the neo-Z(Z): 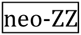 median ratios were ~0.7 in both sexes, significantly lower than those of autosomal controls (AA: 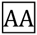) (*p* < 5e-05, Bonferroni-corrected Mann–Whitney U test, Fig 5A). This indicates that overall gene expression on neo-Z is equally reduced in both sexes relative to the inferred ancestral expression. However, the expression reduction was not universal among neo-Z genes, with strongly-expressed genes appearing to be better compensated than more weakly-expressed genes. For example, using *M. sexta* as the reference, the distributions of neo-Z(Z) and 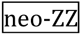 expression are concordant for genes with log_2_(FPKM) values greater than 5 (an arbitrary threshold chosen post-hoc to illustrate this point; Fig 5B). In contrast, below this threshold, neo-Z(Z) expression is shifted significantly towards lower values compared to those of 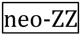 (*p* < 10^−15^, Bonferroni-corrected Komolgorov-Smirnov tests) (Fig 5B). This pattern suggests that in *C. pomonella,* Z~A compensation was local and mostly operated on the most strongly-expressed genes.

**Fig 5.**
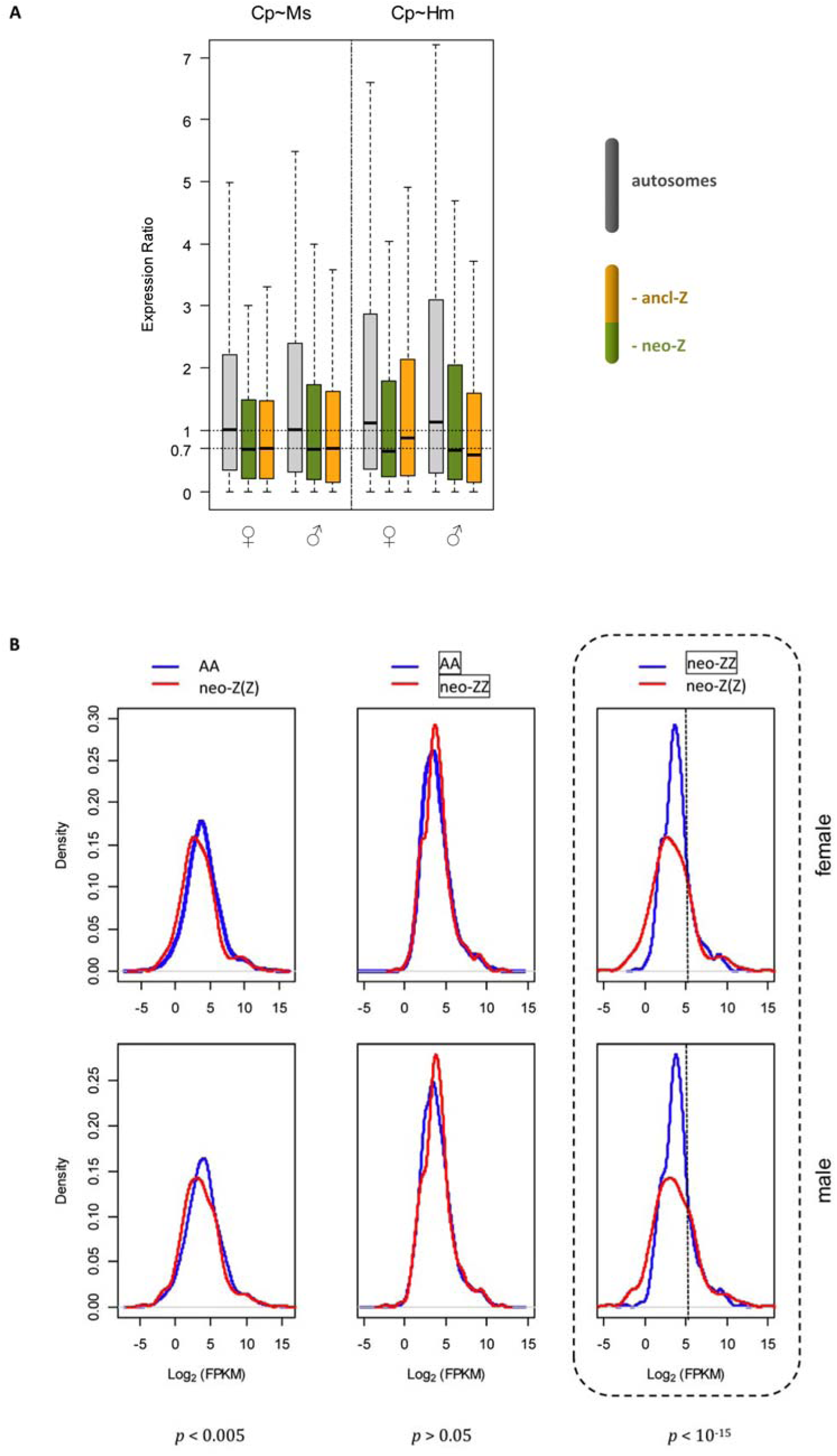
Dosage compensation analysis in *C. pomonella* based on inter-specific data. (A) Expression ratio of orthologs between *C. pomonella* (*Cp*) and reference species (Ms: *M. sexta,* Hm: *H. melpomene*). Grey: AA: 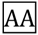, green: neo-Z(Z): 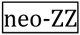 orange: ancl-Z(Z): 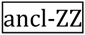. (B) Distributions of expression levels of orthologs between *C. pomonella* and *M. sexta.* Bonferroni corrected *p* values from Komolgorov-Smirnov tests are shown below each category.

The comparative transcriptome method does not infer the ancestral expression level of ancl-Z genes. However, ancl-Z(Z) expression also showed a similar ~30% overall reduction in both sexes compared to *M. sexta.* In comparison with *H. melpomene,* the reduction of ancl-Z expression was unequal between sexes, which corroboratively reflects the minor Z dosage-effect in *H. spp* reported by (Walters, et al. 2015). One possible scenario to explain this reduction in *C. pomonella* ancl-Z expression relative to Z-linked orthologs is that Z-linked expression may have increased among the more derived species. Indeed, an intriguing phylogenetic trend of increasing Z:AA ratios is observed among increasingly derived species, at least for the head, which is the only tissue available for comparison. Specifically, ancl-(Z:AA|ZZ:AA) ratios were the lowest in *C. pomonella* (0.56|0.62), whereas Z:AA|ZZ:AA ratios are 0.63|0.69 in *H. melpomene* (Walters, et al. 2015), 0.761|0.766 in *B. mori* (Walters and Hardcastle 2011) and 0.8|0.83 in *M. sexta* (Smith, et al. 2014). Therefore, the extent of Z~A compensation seems to correlate with the phylogenetic hierarchy in Lepidoptera such that more derived species are generally more compensated. Nevertheless, this taxonomic sampling is sparse and this pattern must be confirmed with data from additional lepidopteran species.

### Exemption from dosage compensation in reproductive tissues

Analyses of dosage compensation (Z~ZZ compensation and Z~AA compensation) on reproductive tissues (ovary, testis and accessory gland) reveal patterns that are distinct from those observed in head and midgut. Considering only genes co-expressed between sexes, which are used to assess Z~ZZ compensation, median autosomal expression was 1.8-fold greater in ovary than in testis (p=0.002, Mann-Whitney U test, Fig 4A). For both ancl-Z and neo-Z genes, the distributions of F:M expression ratios differed significantly (p<0.05, Benjamini-Hochberg-corrected Komolgorov-Smirnov test) from that of the diploid autosomal baseline, corresponding to 1.9- and 1.5- fold dosage effect respectively.

The highly female-biased autosomal expression for co-expressed genes in the gonads is unexpected, which possibly reflects the difference in general transcription intensity associated with oogenesis and spermatogenesis. On the contrary, genome-wide excess of testis-specific genes contribute to a strong male bias in autosomal expression for sex-specific genes in the gonads (see the section below, Fig 6). Interestingly, the opposing bias from these two subsets of genes (co-expressed and sex-specific) resulted in close median values of overall autosomal expression between ovary and testis (Fig 4B). As a reflection of the high divergence, however, overall gene expression between male and female samples was poorly correlated in the gonads (Pearson’s ρ=0.12), compared to the head (Pearson’s ρ=0.92) or the midgut (Pearson’s ρ=0.73).

**Fig 6.**
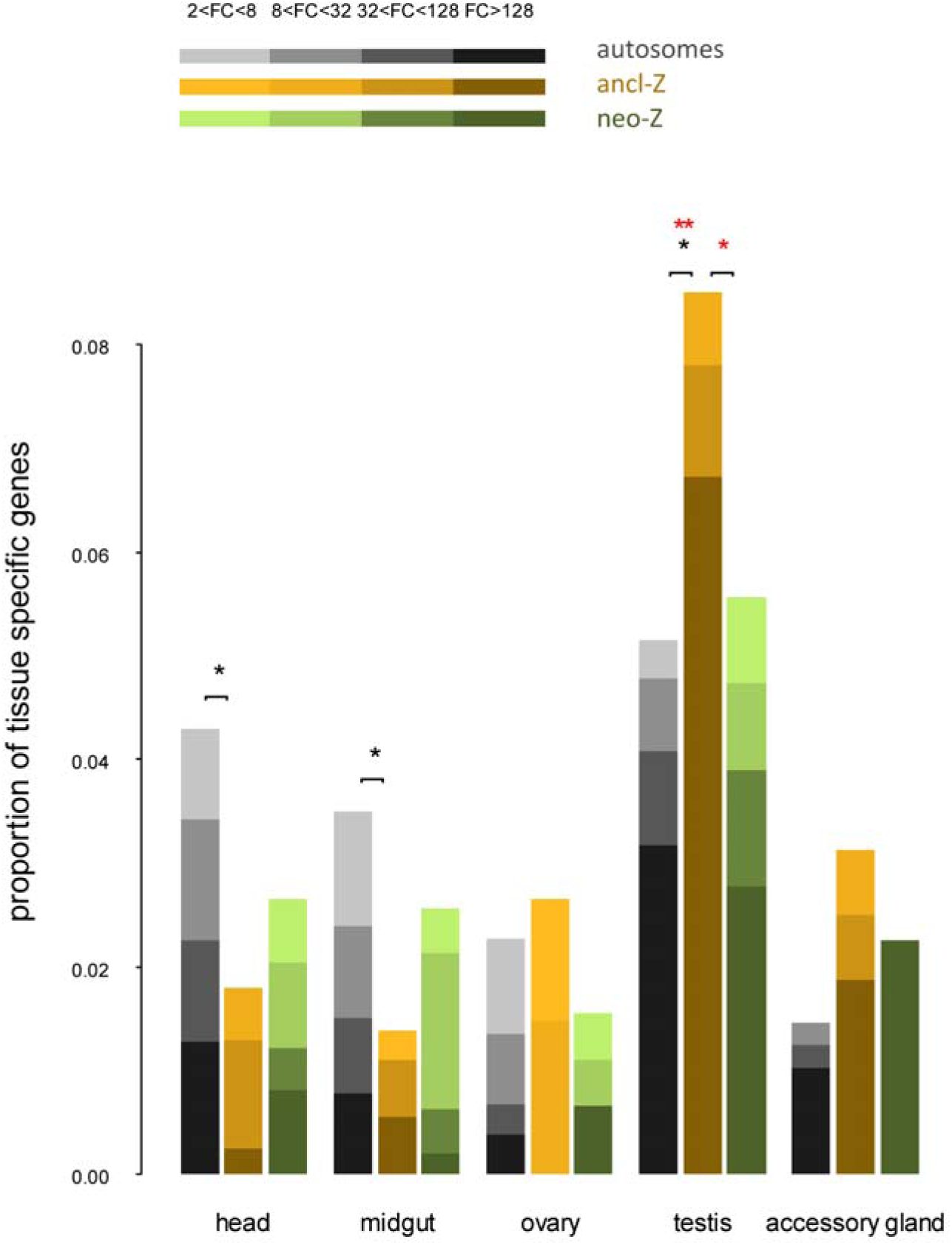
Genomic distribution of tissue-specific genes in *C. pomonella.* Tissue-specific genes were identified under FC cutoff values from 2 to 128. Statistically significant differences assessed by two tailed Fisher’s exact test are indicated on top of bars: * *p*<0.05; ** *p*<0.01 | * FC cutoff = 2; *,** FC cutoff = 128.

For Z~A compensation, neo-Z expression was significantly higher than ancl-Z expression both in the ovary and testis (p<0.01, Bonferroni-corrected Mann–Whitney *U* test, Fig 4B), which is consistent with the pattern in the head and midgut. In contrast, no such difference was observed in the accessory gland (p=1). Among all the tissues, testis and accessory gland had the highest ancl-ZZ:AA and neo-ZZ:AA ratios, which are not significantly different from 1 (bootstrap test, Fig 4C). This result is in good agreement with the *B. mori* data, which also show a median ZZ:AA ratio of 1 in the testis (Walters and Hardcastle 2011). In contrast, ovary had the lowest median ancl-Z:AA ratio (0.48) that did not differ significantly from 0.5 (bootstrap tests, Fig 4C). Although the median neo-Z:AA ratio in the ovary (0.76) was significantly higher than 0.5 (but < 1, bootstrap tests), the distribution of neo-Z expression levels did not differ significantly (p=0.2) from the distribution of computationally-simulated monoallelic autosomal expression levels (S2 Fig, Table S2). It thus appears that in the reproductive tissues, both ancl-Z and neo-Z genes are expressed at levels consistent with underlying chromosome ploidy and there is not a mechanism operating to compensate dosage. This observation contrasts the lack of gene dosage effect between sexes both in the head and midgut, and suggests the exclusion from dosage compensation in the reproductive system.

Additional evidence supporting this conclusion comes from the comparative transcriptome analyses. The expression levels of both 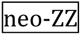 and 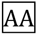 represent gene expression in the unmodified diploid state. The 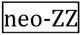: 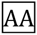 ratios of median expression estimated from head tissues in both reference species were close to 1.1, which were significantly (bootstrap tests) higher than the expected value of 1 (S4 Fig). Strikingly, this value of 1.1 coincides with the neo-ZZ:AA ratios of median expression in *C. pomonella* testis and accessory gland (1.13 and 1.14, respectively), while these ratios were significantly (bootstrap tests) less than 1.1 in all the other tissues (Fig 4B). This unique consistency between species in median expression of neo-Z relative to autosomes suggests that neo-Z gene expression in the male reproductive tissues reflects ancestral expression levels, a pattern that is consistent with the absence of dosage compensation.

### Nonrandom genomic distribution of tissue-specific genes in *Cydia pomonellla:* implications for dosage compensation

Studies on various species, including lepidopterans, have shown that sex chromosomes typically have unusual complements of genes that are expressed predominantly in one sex (sex-biased genes), especially in the germline (Arunkumar, et al. 2009; Khil, et al. 2004; Lercher, et al. 2003; Parisi, et al. 2003; Reinke, et al. 2004; Saifi and Chandra 1999). Walters et al (Arunkumar, et al. 2009) suggested that in *B. mori,* over-representation of strongly-expressed testis-specific genes on the Z chromosome could account for the exceptional ZZ:AA ratio in the testis, yet this explanation was not directly tested. To further address this issue, we explored the interplay between genomic distribution of tissue-specific genes and dosage compensation patterns in *C. pomonella.* We identified tissue-specific genes and normalized the numbers of tissue-specific genes to the total numbers of expressed genes on each chromosomal location (autosomes, ancl-Z and neo-Z segments). Here, tissue-specific genes are defined via differential expression as having consensus up-regulation in a particular tissue as compared to any other tissue for a given fold change (FC) cutoff value. For head and midgut, few differentially expressed genes were identified between the sex-specific samples (20 and 8 for the head and midgut, respectively), which were hence pooled to re-calculate FPKM in order to reduce complexity of the analysis.

Although testis-specific genes were generally overrepresented in the genome compared to other tissue-specific genes, the ancl-Z segment harbored a much larger proportion of testis-specific genes than either the autosomes or the neo-Z segment (Fig 6). This pattern was significant at all FC cutoff values (*p* < 0.05, Fisher’s exact test) and became more pronounced with increasing FC thresholds. Notably, at FC =128 the proportion of testis-specific genes was twice as large on the ancl-Z segment as on the autosomes. In contrast, head- and midgut-specific genes were significantly underrepresented on the ancl-Z segment (*p* < 0.05, Fisher’s exact test). Patterns on the neo-Z for testis-, head- and midgut-specific genes similarly trended in the directions noted for the ancl-Z, but were not statistically significant (*p* > 0.05, Fisher’s exact test). Accessory gland-specific genes tended to be more prevalent on both the ancl-Z and neo-Z segments relative to the autosomes, but the pattern were not statistically significant. However we note that the lack of significance might reflect low statistical power because the total number of genes expressed in the accessory gland *per se* is half as many as those expressed in the head. In addition, we used EdgeR to identify differentially expressed genes under stringent false discovery rate (FDR < 0.05, *p* values for FDR < 0.05), so the number of tissue-specific genes recovered may be an underestimation.

Highly-biased expression is typically associated with high expression levels. Indeed, all the subsets of tissue-specific genes, particularly the testis- and accessory gland-specific genes, were among the most highly-expressed genes in every tissue (S5 Fig). Nevertheless, computational removal of all the 1,247 tissue-specific genes (at FC=2), which took up 12.43% of the entire transcriptome, did not appreciably change the ancl- or neo-Z(Z):AA ratios in any tissue (S6 Fig) compared to the calculation based on the whole data set (Fig 2B). Therefore, it appears that the nonrandom genomic distribution of tissue-specific genes did not confound the assessment of Z~A compensation in *C. pomonella.* In particular, this result lends confidence to the notion that the distinct Z:A ratios reflect largely unmitigated dosage effects and that dosage compensation is absent in reproductive tissues.

Sex-biased gene expression is typically associated with sexual dimorphism and, accordingly, germline-specific genes represent a substantial fraction of sex-biased genes (Ellegren and Parsch 2007). The over-representation of testis-specific genes on the Z chromosome in *C. pomonella* provided further evidence supporting sexually antagonistic selection (Rice 1984). However, this theory cannot explain the under-representation of head- and midgut-specific genes, since these genes do not involve sex-specific expression. We suggest that the lack of general Z~A compensation in the head and midgut would make the Z chromosome a suboptimal environment for genes that require high expression, hence drive selection for their relocation to the autosomes. Consistent with such a scenario, the neo-Z and the ancl-Z segments both showed deficit in head- and midgut-specific genes but to different extent, which corresponds to the differential Z~A compensation between the two chromosomal segments.

### Inconsistencies between *P. interpunctella* and other lepidopteran species may reflect analytical artifacts

Dosage compensation is typically conserved in taxa with highly-conserved sex chromosomes like the Lepidoptera. In that respect, the pattern (Z < ZZ ≈ AA) reported in *P. interpunctella* (Harrison, et al. 2012) seems peculiar since both the more basal species (*C. pomonella*) and the more derived species (*B. mori, M. sexta* and *Heliconius. spp*.) share the same pattern (Z ≈ ZZ < AA). While it may be that *P. interpunctella* represents a true anomaly among Lepidoptera, this inconsistent result motivates a careful consideration of possible artifacts that may offer an explanation for the pattern. Consider first that the analysis of dosage compensation in *P. interpunctella* sampled whole-body adult insects, which contain a substantial fraction of reproductive tissues. Evidence from both *B. mori* (Walters and Hardcastle 2011) and *C. pomonella* point to the exemption of dosage compensation in the reproductive tissues. Thus inclusion of the reproductive tissues is prone to bias the overall assessment of dosage compensation. Second, the FPKM values were not TMM-normalized in the *P. interpunctella* study. As a result presumably, the median autosomal expression is notably higher in the female than in the male. Similarly in the *C. pomonella* data, median autosomal expression was higher in the ovary than in the testis before TMM normalization, while TMM normalization resulted in a distribution of FPKM values in the gonads similar to somatic tissues (S1 Fig). By comparison, TMM normalization did not appreciably change the FPKM values for the head and midgut, which is consistent with the *M. sexta* study where only head was sampled (Smith, et al. 2014). In accord with the rationale of TMM normalization (Robinson and Oshlack 2010), this observation reflects the differences in RNA population between the testis and the ovary relative to the other tissues. Finally, instead of uniformly filtering data across samples, the authors of the *P. interpunctella* study only removed those contigs with below-threshold FPKM values in ‘at least two out of four samples’ (two replicates for each sex). Since the female samples have generally higher FPKM values (without TMM normalization) than the male samples, such filtering procedure is expected to disproportionally remove more contigs with low FPKM values in the male replicates than in the female replicates. Accordingly the compression effect, which as discussed earlier causes disproportionate removal of low FPKM values between the Z-linked and autosomal gene sets, is expected to mask the Z:A expression disparity in the male but not (yet) in the female. Curiously, the arbitrarily-set threshold (FPKM = 4) happens to be the same value under which the Z:A disparity starts to diminish in the *M. sexta* study (Smith, et al. 2014).

Aside from the fact that the raw data from the *P. interpunctella* study were not made publically accessible, clarification of dosage compensation in this species would require sampling tissue(s) in isolation from the reproductive tissues. Furthermore, a broader sampling among lepidopteran lineages will arguably contribute to a definitive answer regarding the conservation of dosage compensation in this order.

## Conclusions

In this study we provide evidence that in the adult soma of lepidopterans Z-linked gene expression is equally reduced relative to the autosomal expression between males and females (Z≈ZZ<AA). This pattern of Z~ZZ compensation coupled with partial Z~A compensation mirrors the pattern of dosage compensation in eutherian mammals. While the mammalian X~XX compensation mechanism (i.e., global X chromosome inactivation in female soma) is long known, recent studies have confirmed that mechanisms of X~A compensation (i.e., up-regulation of the monoploid X chromosome) (Deng, et al. 2013) operate only on a minority of X-linked genes (Chen and Zhang 2015; Julien, et al. 2012; Lin, et al. 2012; Pessia, et al. 2012). In contrast, information on mechanisms of dosage compensation is lacking in Lepidoptera, and only a zinc finger protein (Masc) has been recently identified in *B. mori* (Kiuchi, et al. 2014) and *Ostrinia furnacalis* (Fukui, et al. 2015), which mediates global repression of Z-linked expression during the embryonic stage in both species.

In Lepidoptera, the extent of Z~A compensation seems to vary across species and among tissues, which is in contrast to Z~ZZ compensation that is consistently observed to be complete. In *C. pomonella* in particular, the younger neo-Z genes were subject to greater Z~A compensation than the ancl-Z genes in all tissues examined except for the accessory gland. Strikingly, similar pattern reflecting evolutionary history of X~A compensation is observed in mammals between the ancestral X-linked genes that are conserved on chicken autosomes and newly-acquired X-linked genes (Deng, et al. 2013).

Further, our analyses in *C. pomonella* revealed that the reproductive tissues (ovary, testis and accessory gland) appear to lack dosage compensation that operates in the soma. Exemption from systemic dosage compensation in the germline has been consistently observed in the XX/XY model systems (mammals (Sugimoto and Abe 2007), *Drosophila* (Meiklejohn, et al. 2011) and *Celegans* (Kelly, et al. 2002)). To our best knowledge, however, this is the first report of a purely-somatic tissue (accessory gland) that appears to be exempt from dosage compensation.

In addition, our analyses suggest a possible role of incomplete Z~A compensation in limiting the distribution of head- and midgut-specific genes, but not reproductive-tissue-specific genes, on the Z chromosome. Similarly in *D. melanogaster,* restraints from the dosage compensation mechanism is suggested to account for the general paucity of male-biased genes (Vicoso and Charlesworth 2009) and tissue-biased genes that are not directly associated with reproduction (Mikhaylova and Nurminsky 2011).

The insect order of Lepidoptera is the only female heterogametic (WZ/ZZ) taxon known to date that equalizes sex-linked gene expression between sexes. The tantalizing similarities with regard to patterns of dosage compensation between the two distantly-related groups (lepidopterans vs mammals) with opposing sex chromosome constitutions (WZ/ZZ vs XX/XY) point to a compelling area for future study. To what extent do these taxa share a common molecular mechanism, and to what extent might any shared mechanism be homologous or convergent? The moths and butterflies challenge our traditional view on dosage compensation’s association with sexual heterogamety, while offering a unique opportunity to decipher the evolution of sex chromosome dosage compensation.

## Acknowledgements

We thank David Soderlund and Ping Wang for their valuable comments and input on the manuscript. This research is supported in part by the Federal Formula Fund (6217449) granted to D.K and in part by the Griswold Award for graduate research (granted to L. G.) from the Department of Entomology, Cornell University (1398103-00514).

## Materials and Methods

### Samples, RNASeq and de novo assembly

*C. pomonella* pupae were obtained from a commercial breeder (Benzon Research Inc., Carlisle, PA). The following seven tissue samples were dissected from young adult moths within 24 hours after emergence: head (female), head (male), midgut (female), midgut (male), ovary (female), testis (male) and accessory gland (male). The testis was severed from the rest of the male reproductive tract, which is primarily comprised of accessory gland, along with seminal vesicles, ejaculatory duct and ejaculatory bulb (Fig 1C). In this study, the remaining male reproductive tract is regarded as a single tissue sample and referred to as ‘accessory gland’ for the purpose of simplicity.

Tissues dissected from five individuals were pooled into one sequencing sample, and two sequencing samples (replicates) were used for each tissue sample. RNA was separately extracted from each sequencing sample using the QIAGEN RNeasy Kit (QIAGEN, Foster City, CA), following manufacturer’s instruction and subsequently sent to Boyce Thompson Institute (Ithaca, NY) for the construction of barcoded libraries using standard Illumina library preparation protocols (Meyer and Kircher 2010). The resulting 14 libraries were then pooled into three lanes and sequenced on the Illumina HiSeq2000 platform (150bp paired-end) at the Cornell University Biotechnology Resource Center (Ithaca, NY).

The resulting 150bp paired-end reads (Short Reads Archive accession number XXXX) were first filtered and trimmed using Trimmomatic (V0.32) (Bolger, et al. 2014). The remaining reads from all 14 sequencing samples were pooled for *de novo* assembly on the Trinity platform (Grabherr, et al. 2011; Haas, et al. 2013). The finished assembly contained all ‘Trinity components’ (or contigs) ≥ 200bp in length, which represented the *C. pomonella* transcriptome from the 7 adult tissues.

### Estimation of expressional intensity

Fragments per kilobase per million mappable reads (FPKMs) were used to estimate gene expression intensity. Filtered read sets from each of the 14 sequencing samples were individually mapped back to the transcriptome assembly for FPKM calculation using RSEM (Li and Dewey 2011) contained in the Trinity package. The resulting 14 FPKM sets represented the gene expression in the 7 tissues, each with two replicates. In order to facilitate cross-tissue comparison, FPKM values were further normalized by Trimmed Mean of *M*-values (TMM) method (Robinson and Oshlack 2010). Because the correlation of FPKM values between the two replicates was high for all the tissue samples (Pearson’s ρ ranging 0.84 ~ 0.99, mean = 0.93), average FPKMs were calculated and used in all the subsequent analyses.

### Chromosomal assignment

*C. pomonella* 1:1 orthologous contigs to *B. mori* genes were identified using reciprocal best-hit approach with a BLAST algorithm e-value cut-off of 1E-5. *B. mori* gene sequences and chromosome map with scaffold information were downloaded from SilkDB v2.0 (http://silkdb.org/silkdb). The contigs mapped to *B. mori* chromosome 1 (Z) and chromosome 15 were identified as linked to ancl-Z and neo-Z segments respectively, while those mapping to other chromosome locations were considered to be autosomal genes. Contigs that could not be mapped to a *B. mori* chromosome were excluded from subsequent analyses.

### Comparative transcriptome analysis

RNAseq data from *M. sexta* and *H. melpomene* head samples was accessed from published studies (Smith, et al. 2014) and (Walters, et al. 2015). Both FPKM data sets were TMM normalized within the species and averaged among the replicates. The orthologous pairs between *C. pomonella* and *M. sexta* and their chromosomal locations were identified by mutually mapping to the *B. mori* reference genome. For the *C. pomonella~H. melpomene* comparison, the *H. melpomene* reference genome (http://www.butterflygenome.org/) was used to identify 1:1 orthologs *C. pomonella* and infer their chromosome locations. *H. melpomene* chromosome 11 corresponds to *B. mori* chromosome 15 and the two chromosomes share a high level of snyteny (Heliconius Genome 2012).

We adopted a scaling procedure following (Lin, et al. 2012) to enable comparison of FPKM values between species. Specifically, all FPKM values from one species were linearly adjusted by a common factor in each sex so that the median FPKM values for autosomal genes were the same between the two species to be compared. To avoid infinite values when calculating ratios without changing the medians, all FPKM = 0 values were converted to FPKM = 0.01.

### Differential expression analysis

DE genes were identified using EdgeR (Bioconductor) imbedded in the Trinity package (Robinson, et al. 2010). EdgeR uses false discovery rate (FDR) to determine differentially expressed contigs, which adjusts gene-specific *p* values for multiple tests (Storey and Tibshirani 2003). In order to reduce complexity, for head and midgut, reads from female and male samples were pooled to recalculate for non sex-specific FPKM.

